# Transcriptomic analysis of mitohormesis associated with lifespan extension in *Caenorhabditis elegans*

**DOI:** 10.1101/2025.04.15.648933

**Authors:** Juri Kim, Naibedya Dutta, Gilberto Garcia, Ryo Higuchi-Sanabria

## Abstract

Non-lethal exposure to mitochondrial stress has been shown to have beneficial effects due to activation of signalling pathways, including the mitochondrial unfolded protein response (UPR^mt^). Activation of UPR^mt^ restores function of the mitochondria and improves general health and longevity in multiple model systems, termed mitohormesis. In *C. elegans*, mitohormesis can be accomplished by electron transport chain inhibition, decline in mitochondrial translation, decreased mitochondrial import, and numerous other methods that activate UPR^mt^. However, not all methods that activate UPR^mt^ can promote longevity. These and other studies have started to question whether UPR^MT^ is directly correlated with longevity. Here, we attempt to address this controversy by unravelling the complex molecular regulation of longevity of the nematode under different mitochondrial stressors that induce UPR^mt^ by performing RNA-sequencing to profile transcriptome changes. Using this comprehensive and unbiased approach, we aim to determine whether specific transcriptomic changes can reveal a correlation between UPR^mt^ and longevity. Altogether this study will provide mechanistic insights on mitohormesis and how it correlates with lifespan of *C. elegans*.

## Background & Summary

Often referred to as the powerhouse of the cells, mitochondria play a vital role in cellular respiration and energy production ^1^. However, describing mitochondria as just the “cellular powerhouse” is diminishing their functions and importance considering their numerous other roles such as immune signalling, cellular signalling via reactive oxygen species (ROS) production, calcium signalling, and cell death regulation ^2–5^. Mitochondrial quality control is governed by the regulation of mitochondrial dynamics, which involves a tightly coordinated balance of fission and fusion events ^6^. These processes are critical for maintaining mitochondrial morphology, including their shape, length, and number ^7,8^. Mitochondrial dysfunction is one of the hallmarks of aging as reported by López-Otín *et al*. and it becomes more pronounced during aging ^9^. This progressive decline is characterized by the accumulation of mitochondrial DNA mutations, elevated levels of reactive oxygen species (ROS), and a gradual impairment of the respiratory chain ^10^.

For the last decade, “mitohormesis” has been extensively investigated in the field of mitochondrial and cell biology indicating that sublethal stress on mitochondria ^11^ – such as impairment in electron transfer chain (ETC), inhibition of mitochondrial protein synthesis, decline in mitochondrial membrane potential, alteration in fusion-fission dynamics – triggers an improvement in health and viability of a cell, tissue, or organism and can protect the cells and organisms against more severe stresses ^12–15^. The improvement of health is often ascribed to the activation of a beneficial stress response dedicated to restoring mitochondrial health and function under conditions of stress, which can also have more dramatic organismal outcomes ^16^.

One of the examples of this mitohormetic response is the activation of the mitochondrial unfolded protein response (UPR^mt^), which is a retrograde signal (mitochondria to nucleus communication involving transcriptional changes) mediated by several transcription factors and chromatin remodelling factors ^17^. The most extensively studied cause of UPR^mt^ activation is the failure in mitochondrial protein homeostasis (proteostasis), which arise from the buildup of misfolded proteins, defective mitochondrial translation, imbalance in the ratio of mitochondria-derived to nuclear-derived proteins in the mitochondria, and impairment of mitochondrial protein import ^18^. For example, if ATF5 (ATFS-1 in *C. elegans*), which normally degrades after mitochondrial import, cannot be imported due to a reduction in mitochondrial import under conditions of mitochondrial stress or damage, it instead is imported into the nucleus where it can induce the expression of genes involved in protein refolding, mitochondrial import, and autophagy ^19,20^. As a result, the induction of the UPR^mt^ can promote proteostasis and mitochondrial fitness, which has been shown to extend lifespan in model organisms ^21^. Although the concept of UPR^mt^ was first identified in mammalian systems, it has been most extensively studied in *C. elegans*. The induction of the UPR^mt^ leads to the transcriptional up-regulation of mitochondrial chaperones such as *hsp-6* and *hsp-60*, mitochondrial proteases such as *clpp-4* ^22–24^, and many other genes involved in restoring mitochondrial proteins including mitochondrial import ^21^ and mitophagy ^25^. Importantly, as the molecular mechanisms of UPR^mt^ induction was further elucidated in *C. elegans* and other invertebrates, the existence of a canonical mammalian UPR^mt^ has been challenged. Instead, many studies argue that the mammalian mitochondrial stress response is dictated primarily by the integrated stress response (ISR) ^26^.

In addition to the discrepancy between mammalian and invertebrate mitochondrial stress responses, even studies restricted to *C. elegans* have revealed some controversies. Most importantly, several studies have reported that UPR^mt^ activation is not directly correlated with longevity ^27^. Indeed, there are some studies reporting non-significant changes, or even a decrease in the lifespan of worms with UPR^mt^ activation ^27^. For example, disruption of mitochondrial fusion or fission by RNA interference (RNAi) against *fzo-1* (fusion) or *drp-1* (fission) activates the UPR^mt^, but significantly decreases lifespan ^28,29^. In addition, inhibition of some factors involved in mitochondrial import (*timm-23*, *tomm-40*) extends lifespan of worms while inhibition of other factors also involved in mitochondrial import (*timm-22*) showed no change in lifespan ^30^ or even a decrease in lifespan (*tomm-20*, *tomm-22*) ^21,27^. Among the ETC complexes, only the inhibition of complex II components has been shown to decrease lifespan whereas inhibition of other complexes increased lifespan ^31,32^, despite virtually equal activation of the UPR^mt 33^.

Here, we hypothesize that the overreliance of the *C. elegans* community on single transcriptional reporter as a readout for UPR^mt^ activation may be the cause of these discrepancies. The *hsp-6p::GFP* and *hsp-60p::GFP* reporters are the most widely used transcriptional reporter to assess the activation of UPR^mt^ in *C. elegans* ^24^. Activation of these reporters are almost always abolished when *atfs-1*, encoding one of the central transcriptional regulators of UPR^mt^, was knocked down ^27^. However, the knockdown of *atfs-1* failed to negate the lifespan extension associated with activation of UPR^mt^ in many cases, raising the concern that UPR^mt^ activation is not sufficient for and cannot predict lifespan extension ^27^. However, it is likely that transcriptional reporters such as *hsp-6p::GFP* do not represent the totality of longevity-associated UPR^mt^, highlighting the importance of finding novel genes that correlate better with lifespan extension in UPR^mt^ activating contexts. In pursuit of this, here we selected UPR^mt^ activating conditions by different mitochondrial perturbations, some of which increase lifespan, *nuo-4*, *cyc-1*, *cco-1*, *atp-2;* and *mprs-5* RNAi ^31,32,34^, and some that do not, *mev-1*, *fzo-1* and *tomm-20* ^21,29,35^. We performed an unbiased transcriptome analysis approach (RNA-seq) in an effort to identify gene expression changes that directly correlate with lifespan extension paradigms. We report the genes upregulated upon UPR^mt^ activation in concordance with lifespan extension. Most importantly, we found that canonical markers, like *hsp-6* and *hps-60* are poorly correlated with longevity, thus explaining previous discrepancies in the field. Moreover, by RNA-seq analysis, we demonstrate the downstream transcriptional changes that may underlie the differential effect on longevity between different ETC complex inhibition. Overall, our study provides a comprehensive UPR^mt^ dataset that will be of great utility to evaluate the activation of lifespan extension-associated UPR^mt^ activation and context-specific UPR^mt^ activation.

## Methods

### *C. elegans* culture, collection, and lifespan

Standard RNAi plates for growing *C. elegans* were prepared as previously described ^36^ NGM-RNAi plates: Bacto-Agar (Difco) 2% w/v, Bacto-Peptone 0.25% w/v, NaCl 0.3% w/v, 1 mM CaCl_2_, 5 µg/ml cholesterol, 0.625 mM KH_2_PO_4_ pH 6.0, 1 mM MgSO_4_, 100 µg/mL carbenicillin, 1 mM IPTG. Bacterial cultures were prepared by growing *E. coli* K strain HT115 carrying a pL4440 vector harboring an ampicillin resistance cassette. Vector positive bacteria were grown using LB medium containing 100 µg/mL carbenicillin in a 37 °C shaking incubator for 16 hours to obtain a “saturated” culture. For control, pL4440 empty vector bacteria (ev) were utilized and RNAi conditions use bacteria with pL4440 vectors carrying a sequence targeting the gene of interest, which is under the control of an IPTG-inducible T7 promoter, which allows for bacteria to synthesize double-stranded RNA, which is utilized by the *C. elegans* RNAi machinery for RNAi-mediated knockdown ^31^. RNAi bacteria was mixed 50:50 RNAi:ev to dilute the concentration of RNAi (with matched OD_600_ measurements) to minimize the decrease in animal growth and size that is often found in ETC knockdown conditions. 200 µL of this mixed culture or ev control was seeded onto NGM-RNAi plates 24 h prior to plating worms.

*glp-4(bn2)* temperature sensitive mutant animals were grown at permissive temperature (15 °C), bleached, and L1-arrested for age synchronization ^36^. L1 animals were plated on RNAi plates and control empty vector (ev) plates at 22 °C for 3 days (72 hours) until they reached day 1 adulthood. We opted to use 22 °C for our restrictive temperature as after backcrossing the *glp-4(bn2)* animals 6x to our wild-type N2 lines, we found that animals were fully sterile at 22 °C. Since growth at elevated temperatures, including 25 °C can have effects on animal physiology, we utilized the minimal temperature that displayed sterility. Approximately 800 animals were harvested from all conditions using M9 at the day 1 adulthood for further RNA isolation. Worms were pelleted down by centrifugation (1.2 rcf, 1 min, 25 °C) and the supernatant was aspirated using a vacuum pump. Afterwards, animals were immediately placed into 1 mL Trizol solution and stored at −80 °C until use. Three independent biological replicates were used, which is defined as distinct bleaches of animals from distinct populations (i.e., no shared progeny from a single parent).

*C. elegans* lifespan assays were performed using wild-type N2 lines on standard RNAi plates and were all performed at 20 °C. Animals were synchronized by bleaching and L1 arresting as described above and plated directly onto RNAi plates at the L1 stage. Day 1 adult animals were moved onto FUDR plates (100 µL of 10 mg/mL FUDR were spotted on the bacterial lawn). Animals were scored every other day until all animals were scored as either dead or censored. Animals that exhibit bagging, intestinal leakage out of the vulva, crawling up the side of the plates, burrowing into the agar, or other age-unrelated deaths were censored and removed from quantification. All statistical analysis was performed using Prism7 software using LogRank testing.

### RNAi sequences

All RNAi constructs were isolated from the Ahringer Library ^37^ and the sequence of each RNAi was verified using sanger sequencing (Genewiz) using a universal M13F primer. Verified sequences for each RNAi construct is provided below.

#### atp-2

TAATTTCCGTGTTGCTTGGCGAGTTCTTCGGCCTTCTTGAACACATCATCGATTCCTCCTTGCATGTAGAAAGCAACTTCTGGAAGGTGGTCGAGTTCTCCCTTGAGAATCATGGTGAAACCGCGGATTGTCTCTTCGAGTGATACGAACTTTCCTTGATGTCCAGTGAACACCTCAGCGACTTGGAATGGTTGGGAAAGGAAACGTTGGATCTTGCGGGCACGTGAAACGGTAAGCTTGTCTTCCTCGGACAACTCATCCATACCAAGAATAGCAATAATATCTTGGAGGGACTTGTAATCTTGGAGGATCTTTTGAACTCCACGAGCAATGTCGTAGTGGTTCTGTCCGACAACGTTGGGGTCCATGATACGGGAGGTGGAGTCAAGTGGATCGACAGCTGGGTAGATAGCCAATTCGGCAATACCACGGGACAAGACAGTGGTGGCGTCCAAGTGAGCGAATGTGGTAGCTGGAGCTGGATCAGTCAAATCGTCAGCAGGTACGTAAATAGCCTGGACGGATGTGATGGATCCCTTCTTGGTGGTGGTAATTCTCTCTTGCATACTTCCCATGTCAGTGGCAAGGGTTGGTTGGTATCCGACAGCGGATGGGATACGTCCGAGAAGAGCGGACACTTCAGATCCAGCCTGGGTGAAACGGAAAATGTTGTCAATGAAAAGAAGCACATCTTGTCCTTCCTGGTCACGGAAGTACTCGGCGACAGTGAGACCGGTGAGGCACACACGAGCACGGGCACCTGGTGGCTCGTTCATTTGTCCGTATACGAGAGATACCTTGGAGTTCTTTCCCTTGAGATCAATGACTCCTCCTTCAATCATTTCGTGGTAGAGATCGTTTCCTTCACGGGTACGCTCTCCGACTCCAGCGAACACAGAGTAACCTCCGTGAGCCTTGGCGACGTTGTTAATGAGCTCCATGATAAGCACAGTCTTTCCGACTCCAGCTCCTCCGAAAAGTCCAATCTTTCCTCCCTTGGCGTATGGGGCAAGAAGATCGACGACCTTGATTCCGGTAACAAGAATCTCCTGCTCGACGGACATTTCGACGAATTCTGGAGCTTCAGCGTGGATAGCAGCGAAGTTCTTGGATGCAATTGGACCTCTCTCGTCAATTGGCTCTCCAATGACGTTCATGATACGGCCAAGAGTTTCTGGTCCGACTGGGATCTTAATCGGATCTCCGGTGTCAGCAACTGGCTGTCCACGGACAAGTCCTTCAGTTCCGTCCATAGCAATACATCTGACGACATTGTCTCCGAGATGTTGAGAGACCTCAAGAATGAGTCTTGGGGAACGTCCGACTACCTCGAGACCGTTGAGGATTGGTGGAAGGTTCTCATCGAACTGGACGTCGACGACGGCTCCGATGACAGCGACAATGCGTCCGGAAGCGTTAGCAGCTGTAGCCTTGGCAGAGACCTTGGTGGCAGCGGCAGCTTGCTGGGTGGCAACACCGGTGTGAATTCCCTTCTTGGACTCAACATTGTTTGAGCTGAGGCGGATGGAAGCTGCTGGAAGAGCGCATTTCTGAACGTTGCTTTGAAGGAGACGGGAAGCCGAACGGCTGATAGAGGCTAACGAACGCGAAGCC A

#### cco-1

ATCATCTCCAGCAAGACGAGCGAGAAGCATCTTCTTTTCACGTCCGGTAGCGTGCTCAAGTGGATCTGGGTAGTATCCGTAGTCTTCTGGCGAGGCTTCAGTGGCCAAAGTACGACGGGCCACGGCAGCTGGAGCGACGAGCTTCTTCGAAAGGGCGGCAACAGCCGTCTTAGCAAGTTGAGCCAT

#### cyc-1

##### cyc-1 (8-49)

TTTAGATCCTTGAACCACGGCACGCTGCATTACGTACCACGTCGA

##### cyc-1 (50-208)

TACGAAGCAATGTCGAATGAAGAGAACGGTCCGCTGTGAGCCCATGGTAAAGCGTACGGGTGAACGTTGTCTCCGCTAGCGCTGACCGAGTTCTCCAAAGCGTAAACGAGTCCCATGCCTGAAGCGGCAGTGACTCCGGCAAGGGCCGCCAATCCACG

##### cyc-1 (202-305)

CCTCAGTCATGATTGTGTCAACGAAATGACGGTAATGCAAGAATTTCATCGAGTGGCAAGCTGCGCACACTTGCTTGTATACTTCGTATCCACGTCGTACA

##### cyc-1 (303-555)

ACCTTCACTCCAGCTGGAGCCTCAAGATATCCAGTAAGAAGAGAGAAAACGTAATCATCTCCTCCGTGTCTAGCAAGAGCCATCAATGACAAATCTGGTGGAGCAGCTCCGTTGTTGGCAGCCGCTGCGGCCTTCTTGTTTGGATATGGGTTTGGCAACTTATCGGTCAGCATTCCTGGGCGCTGAATGGATGCTCCCTTATCGTCAACATCATTAATAAGAGCGTCAGCGGCCTCGGCTTTAGCTTCTTCCT

##### cyc-1 (552-748)

CTTAAGAGCCCACTTCTTTCTGGTGTCATGGAATGGCTCTGCGGCCCAGTGCATGAATGCAGAAACATCCTTAGCTTGTTGAGACATTGTTGCTGGTGTTCCGTCCTTGTACTCAATTCCTTCGTCGAACAATTGTTGTGGCATAGAGATGATACCTCCTGGGAAGTATGGATTGTAAGCTTTTCCGTCATCAACCT

##### cyc-1 (744-829)

TTTCTGCGACTTCGTGAATGACCAGATATGTCTCTTTCCGTAGATCAGCACAACAGCCACAAATGGAATCAGAGCAGCGATCTAAA

#### fzo-1

##### fzo-1 (491-1528)

TGATTGGAAAAATTGGACACGCACTGAGTGACGAAAATTCAGATCTTCCGGCAATGGGACAGGATTCTCTTTTGAAAGTGTTTCATCCAAAGAAATCGGAATCAGGGGAATGCCGACTTCTTCAAAACGACGTCGTCATACTTGATAGCCCTGGTGTCGATCTTTCTCCAGAATTCGATAGCTGGATTGACAAACATTGTTTAGATGCAGATGTATTTGTGCTTGTATCAAATGCGGAATCAACATTGACACAAGCCGAAAAGAACTTCTTCCTTCGAGTCGCGAGAAGCTTAGCAAGCCCAACGTATTTATTCTCAATAATCGATGGGACGCCAGTGCAGCAGAAGCAGAAAACATCGAAGATGTAAAAAAGCAGCATTTGACGAGATTCCGTCAGTTTTTGGTCGATGAGCTTGAAGTGTGCTCAGAAAGAGAAGTCAATGATCGAATCTTCTTTGTTTCATCGAGAGAAGTTTTGGAATCACGATTAAAAGCTAGGGGACTGGTACAAAAAGCCTATCAAGCAGAAGGTCATGGCACAAGAGCCTTGGAATTCCAAAATTTCGAACGACATTTTGAGCATTGTATCTCTCGATCGGCGATTCACACGAAATTCGAAGCTCATAACAGGAGAGCTCATGAAATGATCGGAAAAATGCGGCTCAATCTGAATAGCGTGCTAACCTCTGCAGCTGAGCAGAGAAGCAAGCTTCAGAATAATCTAAACGGATCCACCAGGACTTTCAATGAATGCCGTGTAAACTTCACTCAATTTGAAAAAGCTTACCGTGAGCAGACAGAGCAACTGCGAGCCGAAGTTCATCTGAAAGTTTCTGCTGATTTCTTTGAAGAAATAGCACGACTTGATGCAATAATTGATCGATTTGAACAACCTTTCGATGGAAGCTCTTCCGGAATGACAAAGTATAAAGAGGATTTGGCAATCTTTGTGGACAAATGTTTGAGCAGTGATTTGGAAGCTAGATGTACCGGTGGCCTAATGTCGAGAATTTGGAATTTGGAGAATGATATGTTC C

##### fzo-1 (1529-1630)

AATATGTGACAAAAATCCTCGCCGAACCTTATCAGAATAAGCTTGAAGAAGTGTGGCGATATCGTGCTCCTTTCAAATTTAGCATTTGTGTCGATGTCCCTG

#### nuo-4

ATCTCTCATTGCATTGGCAAAAACGAAATCCGAATGTGGAGTACGTTCGAGCACAACTCCTTGACCGGTATTGAGAATATGAGCAAGTGCGTTGAGATATTGATCGAAACGGCAGTTGAAAATGCGATCTTGCATGGCAGCTGAGAGTTCACCACTTGGATTCTTATAGAACATGGAAATGTCTGGAAGACGGTAACGAGCTGGGAATTTATTATAATAATTGCGAAGATCATTTCCGTAGCGATCGACGAGAATATCGTCCATTCTAAATTCTGGGAAATGAACGAAGCCGAGTTGATCAGCAAGTTGTTTGGCTAAAGTTGTCTTTCCGGATCCAATATTTCCTTCAACAACGATAAGCTTGGAATTTTGATGGAAATGAGAACGAGTGTCGTCTTTTAATCCATCAATATAATTGAATCCATTGTGCTTATAGTCCCATGGTTCTGGATGTTCAGATGGAAGTCTTAGAAGACTTTTATGAACAACTCCTCTGACTTGATTTGAGAGTGCTCCTGATCTGGAGCAGGCCAGCAAACCGGTTGCCAGCCCTCCCAATTTGCTCATGGAACCCGGCA

#### mev-1

TAGGCAGTCTTGTTGCTCTTGTTCTGGCAAGAGTTGAAGACAATGGCGAGAGCAAGAATAGCCGAAAGTCCAGATACGAGATATCCCGATTTGTAGATCTGTCCAACATTATTGACTCCCTTAGCCAAATCGAATCCTAAGAAGCGAATTCCGTTAAGAGTATGGAAAATGATGGGGAAAGCAATGATGTACTTGAAGACAGCGGTCACCGCGCATGGTAAGTTCCAGCTACGGATGAAATCCACAAAAGCGGTGAAATCGAACGGCAAAACTGCGAATCCGATTCCTCCGACGAGAAGGGTTCCGGCCATTACACAACCGCTGATTCTATGGAATCCGGAGAGCATCCAGGTCAATTGTGGCTGGTAGACGGTGAGATGTGGAGCGATTGGGCGATTCTTGGAGCGCTGCTTCAACAGGTATTCCCATCCGAACTTCTGGATTGGCGTCTTTGCCTCGGACTTGGTGACGATCGATGTTCCAAAGGAGCGGGAAATCGACGATCTGGCGCCCAAGCGGCACAGAATCGCAGTTGGAATGTTAATCA

#### mrps-5

##### mrps-5 (7-183)

ACTGGATCCAATTCGATAAAATCGATTGAGAGGTCTAACTGGTTGTCTTGTGTTACGACGTCCTTTCTTCTGACCAGATTTTGAAACGGAGGTTAATGTTTTCCACAGTTCTGGACCAGATCTTCTCATAAAGAAGTTCACCGTATTGCTACGGGTCTGGACAAATGGCAAAAGTGA

##### mrps-5 (182-332)

GTATCTCGTTCTTCGAGAATTTTCTTGGTTCCACCCATTGAATCTCGAATTTCATCTTCCGTCTGTTCAGCAATTGACATCAAATTTTGATTTTCTGTCTCTCTCATTCTGATTGGAGCATTCAGGCCAGCAAATTCGATTTTCATTGGAC

##### mrps-5 (330-769)

TCAGTCTGGGATGGCATGTAAGACCGAATCCACGTGGGCGACGTTGTGCAAATACTCGAGTGTTACGGCATTCAGCATAAAAGTCCTGATAAATCGTACGTCCTTCATGAAGTTCAACGTGGAATAGCTTTCTGGATGCCATTCCCATTCCATTAATGATAGCTGTAGTGGTACGATGAATTGGTGCTTTTCCAACAGCATATCCAGCAAGTCCTCGT CCATTTCCAGTAACAACTAGTGCTGACATTGTGTGAACTCTTCCAAATACATTTGTCATGTTAGAAGTACGCTTAACCTCTAAACAATAAGTTTCAAAATCATCGAAATTCACACCATCGAGTGGTGGTGGTGCACCAAGTTTCTGTCCGACAAGTTGTGTTCCAGAGAATCCACGCTCCATCGGATGCAGTTTTTCACGGTTGCGCTTCTTTTTTCCAC TA

#### tomm-20

##### tomm-20 (1-144)

GAATGTCGGACACAATTCTTGGTTTCAACAAATCAAACGTCGTTTTGGCTGCTGGAATTGCTGGAGCCGCTTTCCTCGGCTACTGCATTTACTTCGATCATAAGAGAATCAACGCTCCAGACTACAAGGACAAGATTAGGCAAA

##### tomm-20 (145-437)

AGAGACGTGCCCAGGCTGGAGCAGGAGGAATGGCTCCACGTCGTCCAGCTGCAGCAGGCAATGATGCTGCTCCAGATGTCACCGATCCATCCCAAATGCAAAGGTTCTTCTTGCAAGAGGTTCAACTCGGAGAAGAACTCATGGCAGCCGGAAATGTTGATGAAGGTGCCGTGCACATTGCCAACGCAGTCATGCTCTGCGGTGAATCTCAACAACTCCTGTCTATCTTCCAACAAACATTGTCGGAAGATCAGTTCCGTGCTGTTGTTCAACAATTGCCGTCGACTCGCGAA

##### tomm-20 (438-633)

CGTCTTGCGGAAATGTTCGGAGCGAAAGCAGATGAGGCAGAAAACGAGCCACCAATGGTTCAATACCTCGGAGACGGACCTCCACCAGCACAAATCCAAGAGCTTATCGATGACACCGACGACTTGGAGTAATGGATAATGATTTAAAAGTTTTATTAACTTTCAATTTAATTCTTCGTATTTTTCGTTCACCAAC

### RNA isolation

For RNA isolation, animals were flash-frozen with liquid nitrogen and thawed in a 37 °C bead bath three times with 30 s vortexing in between. After the final thaw the samples were mixed with 200 μL of chloroform (5:1 trizol:chloroform ratio) and vortexed for 30 s before being transferred to a heavy gel phase-lock tube (VWR, 10847–802) for the aqueous separation of RNA by centrifugation (15,871 rcf, 10 min, 4 °C). The upper aqueous phase was moved to an Extracta Plus DNA Removal Column, centrifuged, mixed 1:1 with isopropanol, then the total RNA purification was performed using a Quantabio Extracta Plus RNA Kit (95214-050) according to the manufacturer’s directions.

### RNA-Seq library preparation and sequencing

RNA quality control, library preparation, and RNA-sequencing was performed at Novogene using their standard pipeline. Sample purity, quantification, and integrity is measured using an Agilent 5400 system. A table of RNA integrity number (RIN) are available in **Table S1**. RNA library preparation includes poly(A) capture to enrich for mRNA and cDNA reverse transcription. mRNA sequencing is performed using NovaSeq25B flow cell on an Illumina PE150 with 20 million paired reads.

### Bioinformatic analysis

#### Adapter trimming and quality control

The raw pair-end RNA reads were trimmed with their adapters using TrimGalore version 0.6.10 argument of FastQC ^38^ with the following parameters: “trim_galore --fastqc_args “--outdir TrimGalore/PostTrimQC/” --illumina --stringency 3 --o TrimGalore/ --cores 6 --paired *.fq.gz”. Then the raw reads were checked with their quality by Fastqc version 0.11.9 using following parameters: “fastqc -o PostTrimQC --threads 20 *.fq.gz”. The fastqc results were summarized using Multiqc version 1.12 as shown in **Fig. 2**.

#### Read alignments and readcount generation

Trimmed reads were mapped to *C. elegans* reference genome (WBcel235) ^39^ using STAR version 2.7.10 ^40^ with the following parameters: “STAR --genomeDir $GENIN --readFilesIn $f $f2 --readFilesCommand zcat --runThreadN 20 --outFilterMultimapNmax 200 --outFilterIntronMotifs RemoveNoncanonicalUnannotated --alignEndsProtrude 10 ConcordantPair --limitGenomeGenerateRAM 100000000000 --outSAMtype BAM SortedByCoordinate --outFileNamePrefix $outname”. To determine the number of reads mapped to the exon with the C. elegans reference genome annotation, featureCounts function of Subread version 2.0.3 ^41^ was used with the following parameters: “featureCounts -D 1500 -p --countReadPairs –primary -T 20 -s 0 -a /home/sanabriaseq/Genomes/Celegans/Caenorhabditis_elegans.WBcel235.107.gtf -o 073124_UPRmt_featureCounts.txt *.bam”.

#### Differential gene expression analysis and transcript read correlation test

The summarized featureCount readcounts were imported into R version 4.3.1 and the differential gene analysis was performed using the DESeq2 version 1.32.1 ^42^ using RNAi condition as a variable. Differentially expressed genes (DEGs) were defined by adjusted p-value < 0.05. Correlation of gene expression across all samples was evaluated using the pairwise Spearman rank correlations. Principal component analysis was performed on the variance-stabilized count matrices to assess the overall separation of samples across various RNAi conditions.

#### Gene Ontology (GO) analysis

To explore the biological pathways that were significantly changed in each UPR^mt^ activating condition and pathways that were altered in lifespan extending conditions, GO analysis was performed using the package clusterProfiler version 4.10.1 ^43^ in R version 4.3.1 with org.Ce.eg.db as an organism. The Benjamini-Hochberg q-value cutoff of 0.05 was used unless indicated otherwise in the Figure legends.

#### Cross comparison of UPRmt transcriptomic profiles

The DEGs identified in this study were compared with the DEGs of UPRmt induction from publicly available dataset. Both the genesets from C. elegans ^44^ and human cell ^44,45^ were compared to the DEG set from this study. The Upset plots were plotted using the R package UpSetR version 1.4.0 ^46^. The human orthologs of the *C. elegans* genes were determined using OrthoList 2.0 ^47^. The normalized readcounts of the 283T cells upon the treatments of doxycycline, FCCP, MitoBloCK-6 and actinonin were fitted to the linear model using the lmFit() function of R package limma version 3.62.2 and eBayes() function of limma to shrink the variance toward the global value ^48^.

### Data Records

The raw data is deposited in Annotare E-MTAB-15031.

### Technical Validation

We extracted RNA from at least 800 worms per condition to ensure adequate RNA quantity and quality from each biological replicate, using three biological groups per condition (**Fig. 1**). Of note, to eliminate the interference of the germline genetic information, we used *glp-4(bn2)* animals which is a temperature sensitive germline proliferative mutant ^49^. After the library construction, we performed FastQC analysis on each RNA sequencing library with MultiQC. All the libraries showed approximately similar Phred scores >= 37, indicating that there was no systemic bias favouring any condition (**Fig. 2a**). Phred score represents the quality of nucleotide identification from sequencing data and higher Phred scores indicate higher confidence in the base call ^50^, suggesting our RNA sequencing libraries were generated with sufficiently high qualities. Then we mapped the RNA sequencing libraries to the *C. elegans* reference genome WBcel235 using the STAR tool. The overall mapped unique and duplicated reads were sufficient and consistent across all samples (**Fig. 2b**).

**Figure 1.**
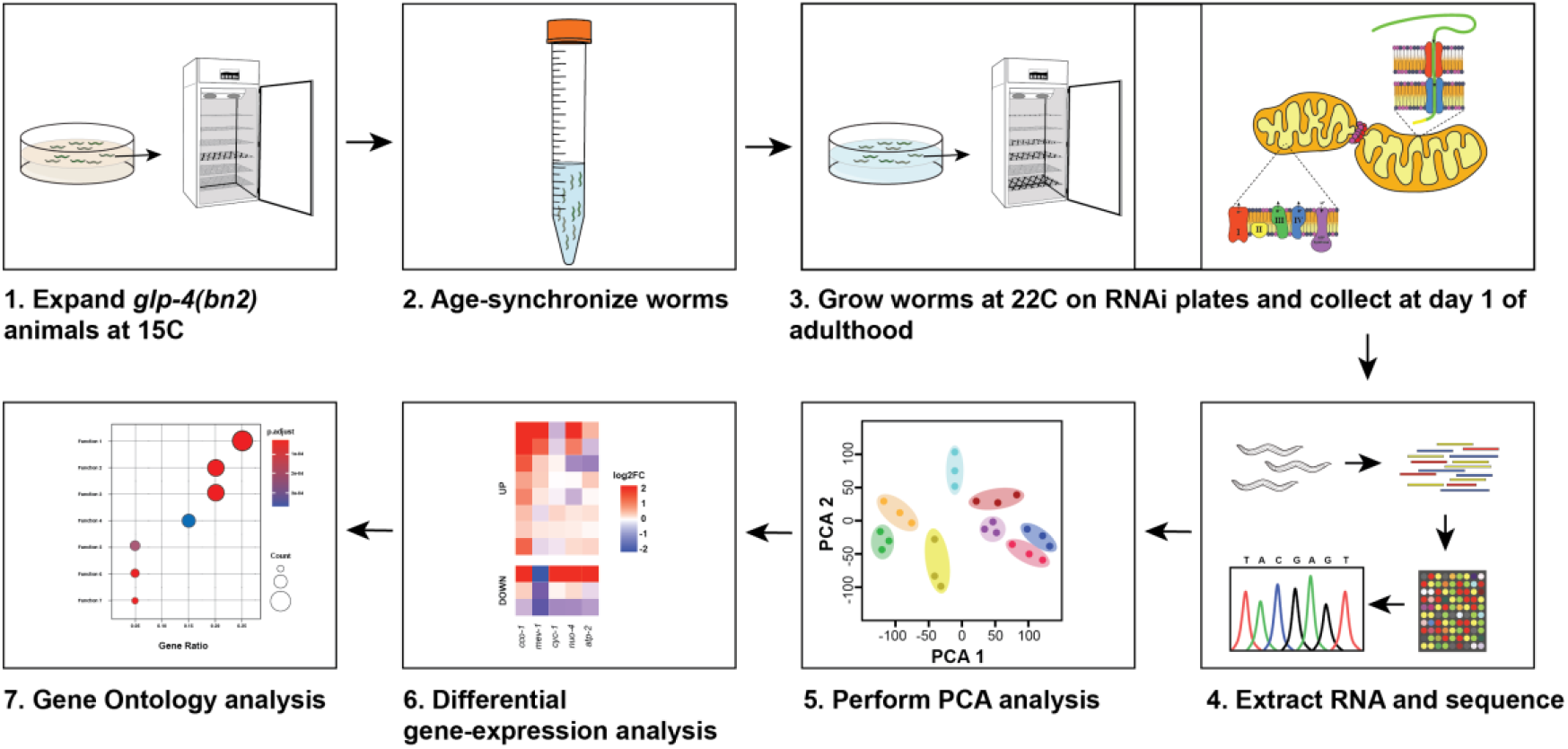
Experimental design and analytic pipeline. *glp-4(bn2)* animals were expanded at 15 °C, the permissive temperature. The animals were age-synchronized by a standard bleaching method followed by L1 arresting for tighter synchronization. Animals were grown on each RNAi bacterial diluted to 50% with empty vector (ev) RNAi to minimize the growth-defects observed in many mitochondrial stress conditions. Animals were exposed to RNAi or an ev control starting from the L1 stage and grown at 22 °C for 72 hours. Day 1 animals were collected, RNA was isolated using a QuantaBio RNA Extraction Kit, and sequenced by Novogene. After sequencing, reads were mapped to the *C. elegans* reference genome WBcel235 and counted. The unique or duplicated reads were compared for each library, and the gene expression of the groups were compared using PCA analysis. Differential gene expression was performed by DESeq 2, and gene ontology analysis was run with clusterProfiler.

**Figure 2.**
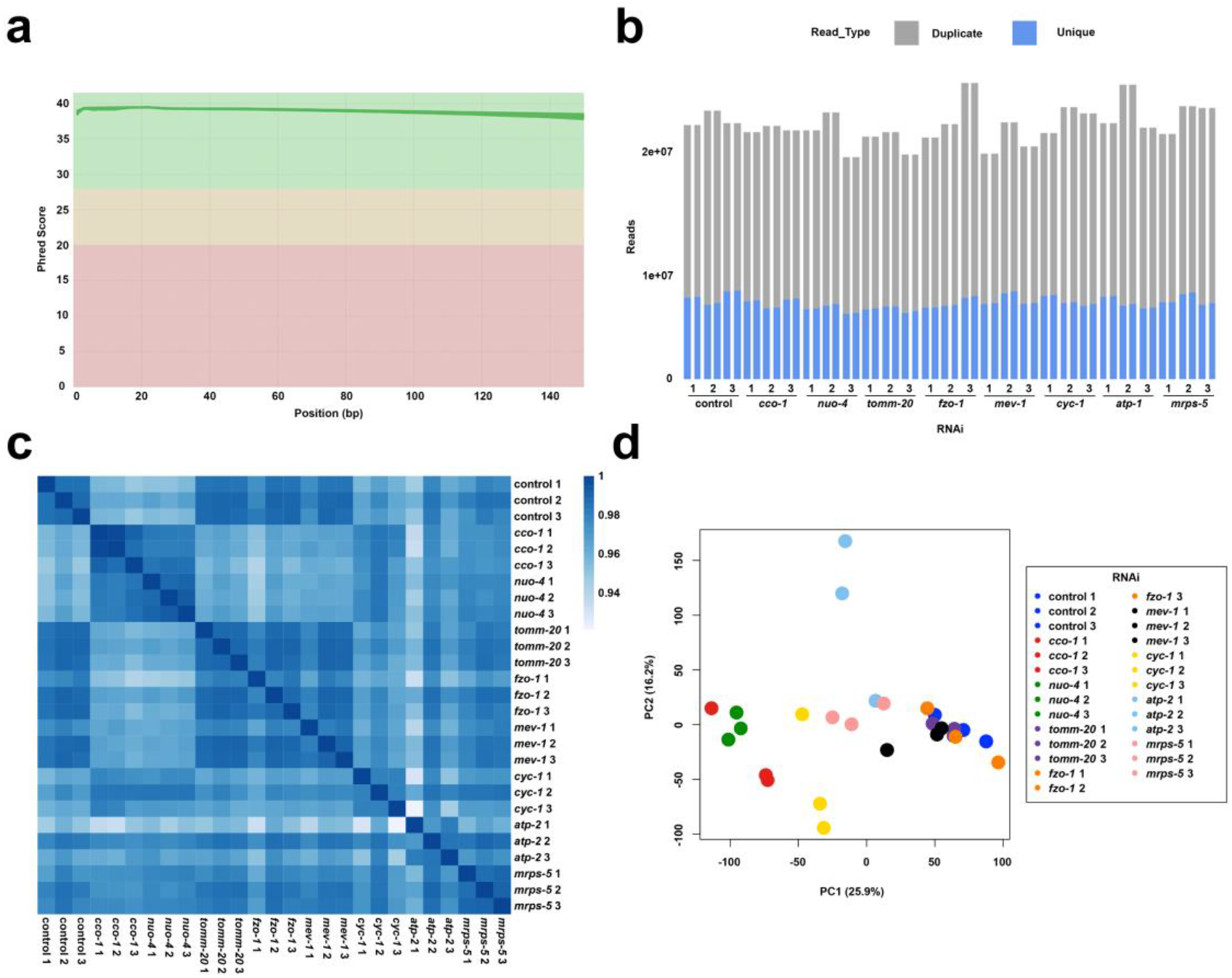
Quality control for RNA-seq libraries of UPR^mt^ activation in *C. elegans* exposed to mitochondrial stress. (**a**) Mean quality score (Phred score) of each RNA sequencing library. X-axis and Y-axis indicate the base pair position of each sequence and the Phred score, respectively. The graph was generated by MultiQC tool (ref). (**b**) The number of unique (blue) and duplicated (grey) reads from each pair-wise sequencing library (n=3). (**c**) Spearman correlation plot of all RNA sequencing libraries. (**d**) PCA plots of UPR^mt^ activated *C. elegans*.

The read counts generated by featureCount were carefully filtered using the iteration algorithm in R (see **Code Availability**) and the unwanted variation resulting from any other factor other than the difference in the sample condition was corrected using SVAseq package version 3.50.0 in R ^51^. The surrogate variable of 2 was used to eliminate the minimal unwanted variation; surrogate variable higher than 2 was avoided in order to prevent extreme artificial correction, which was carefully assessed by examining correlation plot, multi-dimensional scoring plot, and principal component analysis plot. The RIN values (**Table S1**) of the RNA were incorporated into the null model to exclude its influence in variance, whereas the variation resulting from the different RNAi condition was incorporated into the test model of SVAseq. Finally, the biological replicates were used as a variable in the batch correction while performing DESeq analysis.

Each biological replicate shared similar expression profiles as indicated by similar Spearman correlation (**Fig. 2c**) and close clustering using principal component analysis (**Fig. 2d**). *cco-1* and *nuo-4* RNAi conditions clustered most closely to each other, which was expected as both conditions involve RNAi knockdown of an ETC component, which results in a significant lifespan extension. *cco-1* RNAi lifespan extension has been previously reported ^52^ and we confirmed lifespan extension by *nuo-4* RNAi (**Fig. S1**). Interestingly, RNAi knockdown of *tomm-20*, *fzo-1*, and *mev-1* also clustered closely, despite their distinct roles in mitochondrial import, mitochondrial fusion, and ETC complex II activity. However, these three conditions were those that did not directly influence longevity ^21,29,35^, suggesting that transcriptional response of UPR^mt^ that does not correlate with longevity are more similar to each other than those that do (**Fig. 2c-d**). Overall, the robust clustering of biological replicates and physiologically similar conditions confirm the validity of our overall RNA sequencing results.

## Results

Previous literature has identified that the inhibition of *nuo-4*, *cyc-1*, *cco-1*, and *atp-5* which are components of ETC I, III, IV, and V (ATPase) extended the lifespan of *C. elegans* while the inhibition of *mev-1* (ETC II component), *fzo-1* (mitochondrial fusion protein), and *tomm-20* (mitochondrial membrane protein) had either no impact or decreased lifespan ^21,29,35^. To determine whether a specific transcriptomic signature exists for UPR^mt^ activation associated with lifespan extension in *C. elegans*, the shared DEGs among *nuo-4*, *cyc-1*, *cco-1*, and *atp-5* knockdown (collectively referred to as long-lived UPR^mt^) were compared with the DEGs of *fzo-1*, *mev-1*, and *tomm-20* knockdown (collectively referred to as short-lived UPR^mt^). There were 525 commonly upregulated DEGs of long-lived UPR^mt^ animals that were not significantly upregulated in short-lived UPR^mt^ animals (blue area in **Fig. 3a**), and 234 commonly downregulated DEGs of long-lived animals that were not significantly downregulated in the rest (blue area in **Fig. 3b**).

**Figure 3.**
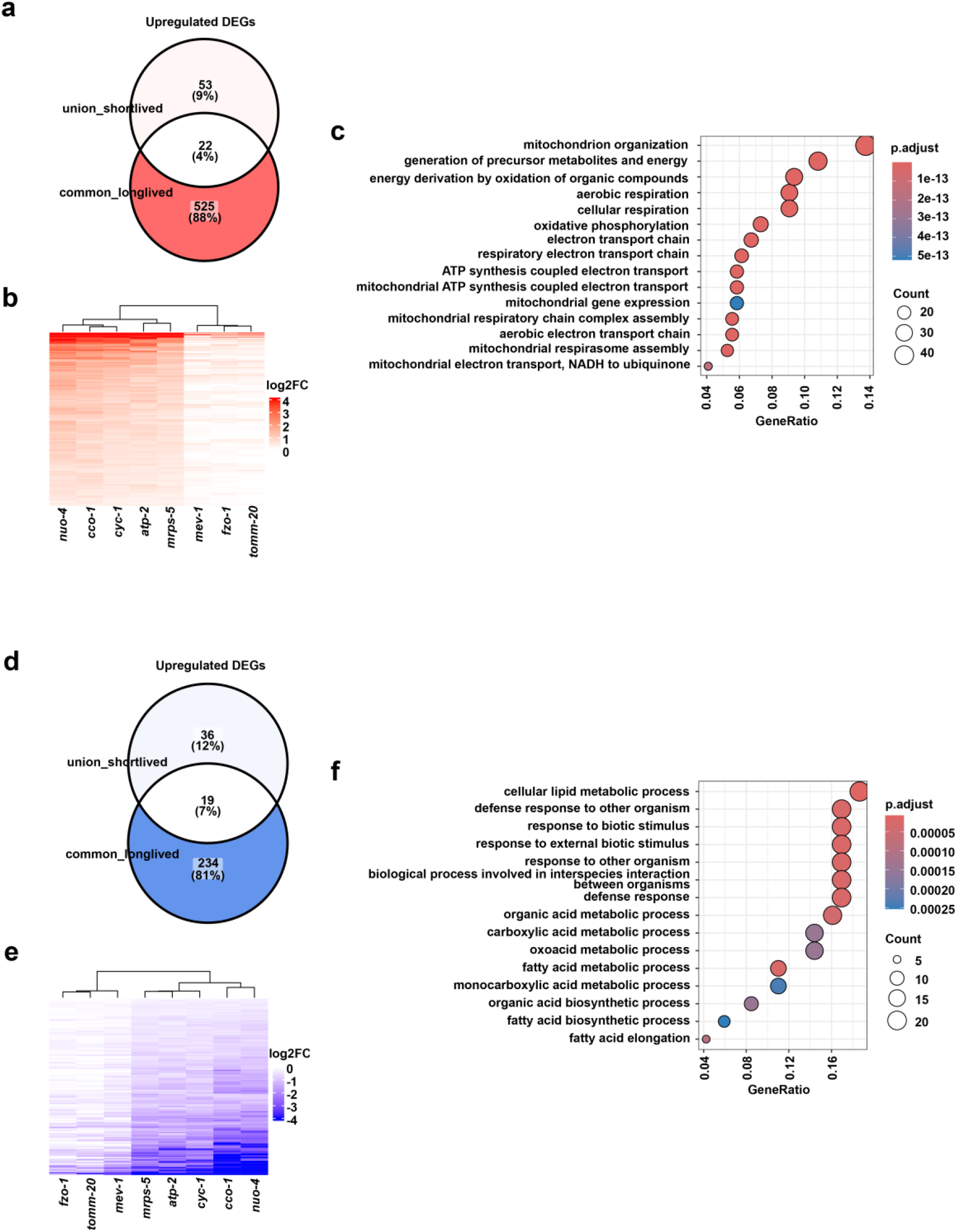
Comparison of gene expression of UPR^mt^ activation in long-lived and short-lived conditions. (**a**) Venn diagram of genes 1) significantly upregulated in all five long lived groups (*atp-1* RNAi, *cco-1* RNAi, *cyc-1* RNAi, *nuo-4* RNAi, *mrps-5* RNAi); *common_longlived*, and 2) significantly upregulated in at least one short lived group (*fzo-1* RNAi, *mev-1* RNAi, *tomm-20* RNAi); union_shortlived. The blue area indicates the genes that are commonly upregulated in long lived groups but not in short lived groups. (**b**) Venn diagram of genes 1) significantly downregulated in all five long lived groups; *common_longlived*, and 2) significantly downregulated in at least one short lived group; union_shortlived. The blue area indicates the genes that are commonly downregulated in long lived groups but not in short lived groups. (**c**, left) GO analysis of the genes that are upregulated in all long-lived groups but not in short-lived groups. (**c**, right) Heatmap of upregulated DEGs from *common_longlived* animals. (**d**, left) GO analysis of the genes that are downregulated in all long-lived groups but not in short-lived groups. (**d**, right) Heatmap of downregulated DEGs from common_longlived animals. All DEGs were selected for adj p-val < 0.05, and the GOs were biological processes (BP) with q-value < 0.05 in **a** and **b**.

To gain a better understanding of the biological processes associated with long-lived UPR^mt^, GO analysis was performed on the upregulated or downregulated DEGs only in the long-lived conditions. GO analysis revealed that long-lived UPR^mt^ was enriched with genes associated with mitochondrial organization, peptide (amino acid) metabolism, and aerobic respiration for upregulated genes and lipid metabolism and innate immune response for downregulated genes (**Fig. 3c** and **3d**). The extended DEG list is provided in **Table S2**.

To understand the unique GO terms of each individual UPR^mt^ condition, GO analysis was performed on up- or downregulated genes unique to each specific condition (**Fig. 4a**). *atp-1* knockdown was enriched with upregulation of genes associated with ribosome biogenesis, translational initiation, translation, and proteasome complex, and downregulation of genes associated with actin cytoskeleton organization, muscle development and contraction, and the vacuole. Both *atp-1* and *cco-1* inhibition was enriched with upregulated genes associated with the ribonucleoprotein granule. *cyc-1* inhibition was enriched with upregulation of genes associated with transmembrane transporter activity, and downregulation of DNA replication, mRNA splicing regulation, and programmed cell death. *nuo-1* knockdown was enriched with downregulation of genes involved in exo-/endocytosis and cytoplasmic/nuclear exosome. Finally, *mev-1* knockdown showed upregulation of genes associated with the vacuolar membrane and membrane transporter activity and downregulated with tricarboxylic acid (TCA) cycle, ETC, response to oxidative stress, and heme binding.

**Figure 4.**
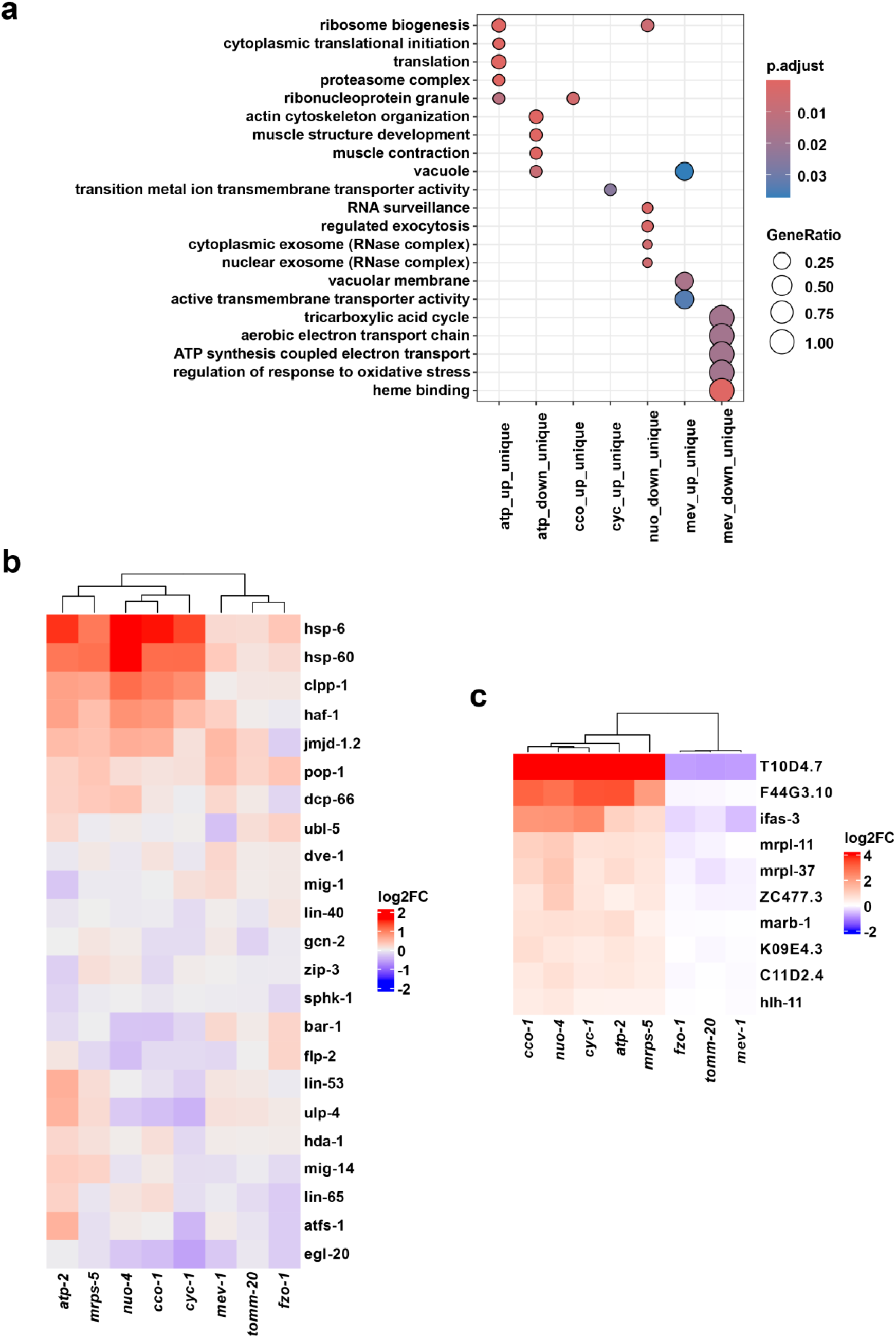
Identification of a novel UPR^mt^ signature associated with longevity. (**a**) GO analysis of uniquely up- or downregulated DEGs from each RNAi condition not shared with another condition. All DEGs were selected for adj p-val < 0.05, and the GOs were biological processes (BP), molecular functions (MF), and cellular components (CC) with q-value < 0.5. (**b**) Heat map of log2FC of the genes known associated with UPR^mt^ listed in AmiGO2. (**e**) Heat map of log2FC of genes significantly upregulated in all long-lived RNAi knockdown conditions (*cco-1, nuo-4, cyc-1, atp-2, mrps-5*), but either unchanged or downregulated in all RNAi knockdown conditions that do not display a lifespan extension (*fzo-1, tomm-20, mev-1*).

Next, to determine gene expression change associated with a “canonical” UPR^mt^ signature in each condition, UPR^mt^ genes were extracted from the database AmiGO 2 (https://amigo.geneontology.org/amigo) and the heatmap of log_2_FC of UPR^mt^-related genes was generated. Interestingly, none of these genes showed uniquely similar gene expression changes across all long-lived UPR^mt^. The only genes that showed common changes across all long-lived UPR^mt^ conditions displayed similar gene expression changes in short-lived UPR^mt^ conditions (**Fig. 4b**). Specifically, *hsp-6* and *hsp-60*, the two genes most commonly used as single-gene reporters for UPR^mt^ induction, displayed a significant increase in expression across all long-lived and short-lived conditions. The overreliance on these two reporters could potentially explain why it has been difficult to predict the longevity of *C. elegans* using the activation of UPR^mt^, since *hsp-6* and *hsp-60* are not unique to long-lived UPR^mt^, but rather is a robust marker for general UPR^mt^. To this end, we generated a gene list that better correlated to the long-lived UPR^mt^ activation with either downregulation or no change in short-lived UPR^mt^ (**Fig. 4c**). Usage of these 10 genes, especially *T10D4*.*7* and *F44G3*.*10*, which display a dramatic increase in expression in long-lived UPR^mt^ conditions and downregulation or no change in short-lived UPR^mt^, respectively, are likely to be more reliable to correlate UPR^mt^ activation with longevity effects.

Finally, to determine whether RNAi knockdown of ETC components share general changes to transcriptome signatures, DEG analysis and GO analysis were performed within the five ETC knockdown conditions. GO analysis of the genes significantly up-/downregulated in *nuo-1* (complex I component), *mev-1* (complex II component), *cyc-1* (complex III component), *cco-1* (complex IV component), and *atp-2* (complex V or ATPase component), but not in other conditions, was performed and revealed that lipid metabolism, more specifically acyl-CoA biosynthesis, is significantly downregulated (**Fig. 5a**). There were only 5 commonly upregulated DEGs and 5 commonly downregulated DEGs across all ETC complex knockdowns, suggesting inhibition of distinct ETC subunits have surprisingly unique effects on gene expression (**Fig. 5b**). Most of those genes were unknown except for the two genes: *dhs-3* and *lpr-3*. *dhs-3* is involved in response to oxidative stress and expressed in lipid droplet ^53^ and *lpr-3* encodes lipocalin that is required for ECM remodelling and organization ^54^.

**Figure 5.**
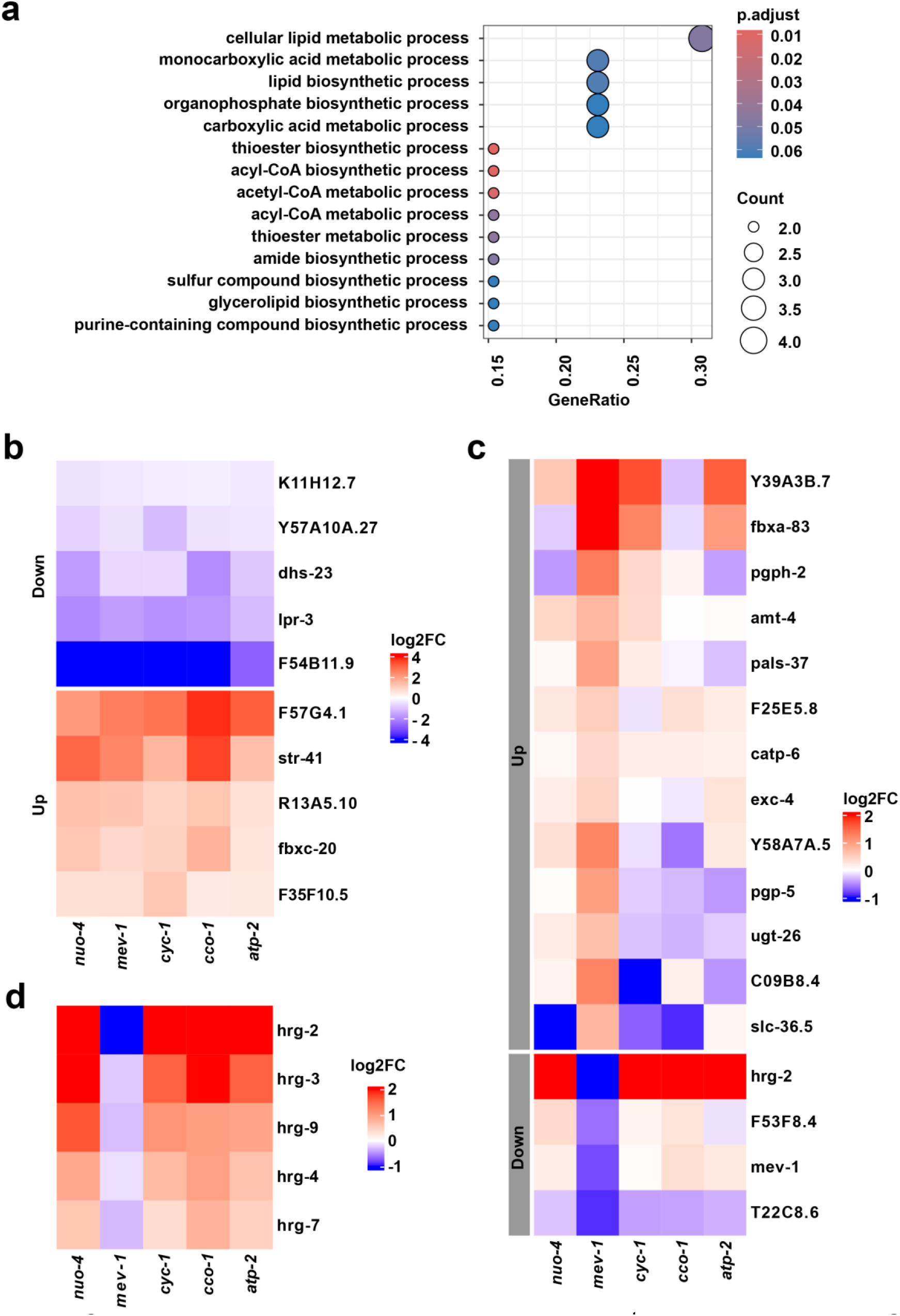
Comparison of gene expression among UPR^mt^ conditions upon ETC inhibition. (**a**) GO analysis of DEGs commonly downregulated upon RNAi knockdown of genes encoding ETC proteins (*nuo-4*, *mev-1*, *cyc-1*, *cco-1*, and *atp-2*. Benjamini Hochberg P-value cutoff=0.07 was used. (**b**) Heatmap of DEGs up- or downregulated <u>only</u> in ETC complex knockdown groups. (**c**) Heatmap of DEGs up- or downregulated <u>only</u> in *mev-1* RNAi group but not in other ETC complex RNAi groups. (**d**) Heatmap of hrg genes upon ETC complex knockdowns.

ETC complex II has historically been shown to be distinct from the other complexes. While complex I receives electrons from NADH, the electron source of complex II is FADH_2_ ^55^. Also, complex II directly oxidizes succinate to fumarate in the TCA cycle, playing a dual role in TCA cycle and electron transfer ^56^. Furthermore, unlike complex I, III, and IV, complex II does not pump protons across the mitochondrial inner membrane ^55^. In an effort to determine the difference between complex II inhibition versus the other complexes, genes that are significantly up-/downregulated specifically in *mev-1* knockdown were visualized (adj p-value < 0.05) in the heatmap in **Fig. 5c**. Interestingly, a majority of these genes do not show consistent changes across the other 4 long-lived ETC knockdown conditions, with the exception of *hrg-2*. *hrg-2* encodes a type I membrane protein that binds heme and is strongly downregulated in *mev-1* knockdown and highly upregulated in the other 4 conditions. Then, we further investigated the expression levels of all the heme responsive genes (hrg) that were significantly upregulated in four ETC knockdown conditions (*nuo-4*, *cyc-1*, *cco-1*, and *atp-2* RNAi) (**Fig. 5d**). Interestingly, all the hrg genes that were differentially expressed in our experiment—*hrg-2, -3, -4, -7*, and *-9* were upregulated in *nuo-4*, *cyc-1*, *cco-1*, and *atp-2* RNAi conditions and downregulated in *mev-1* RNAi condition. The HRGs of *C. elegans* play crucial roles in maintaining the heme homeostasis, *hrg-2* being expressed in hypodermis and *hrg-3, -4, -7*, and *-9* expressed in intestine. They bind to heme and transport it between organs, facilitating the heme utilization. This suggests that the difference in longevity effects of complex II inhibition versus the other ETC complexes could potentially result from the failure to activate hrg genes, though further study to elucidate the mechanism is required.

Next, to display some examples of how our dataset can be utilized further, we compared our DEG sets of UPR^mt^ induction with genesets from several different publicly available data associated with mitochondrial stress. The DEGs from our *nuo-4* (ETC complex I component) RNAi condition (adj p-value < 0.05) was first compared with the *C. elegans* UPR^mt^ genes induced by the treatments of mycothiazole (MTZ), 8-O-acetylmycothiazole (8-OAc), and rotenone, which are all chemical inhibitors of ETC complex I function (adj p-value <0.05) ^44^. There were 122 overlapping DEGs across the chemical treated and *nuo-4* RNAi treated conditions. *nuo-4* RNAi treated worms shared 246, 27, and 6 DEGs with MTZ treated, Rot treated, and 8-OAc treated worms, respectively. *nuo-4* RNAi, MTZ, and Rot treated worms shared 117 DEGs (**Fig. 6a**). Thus, while differences certainly exist between genetic or chemical targeting of complex I, there are definitely overlapping DEGs.

**Figure 6.**
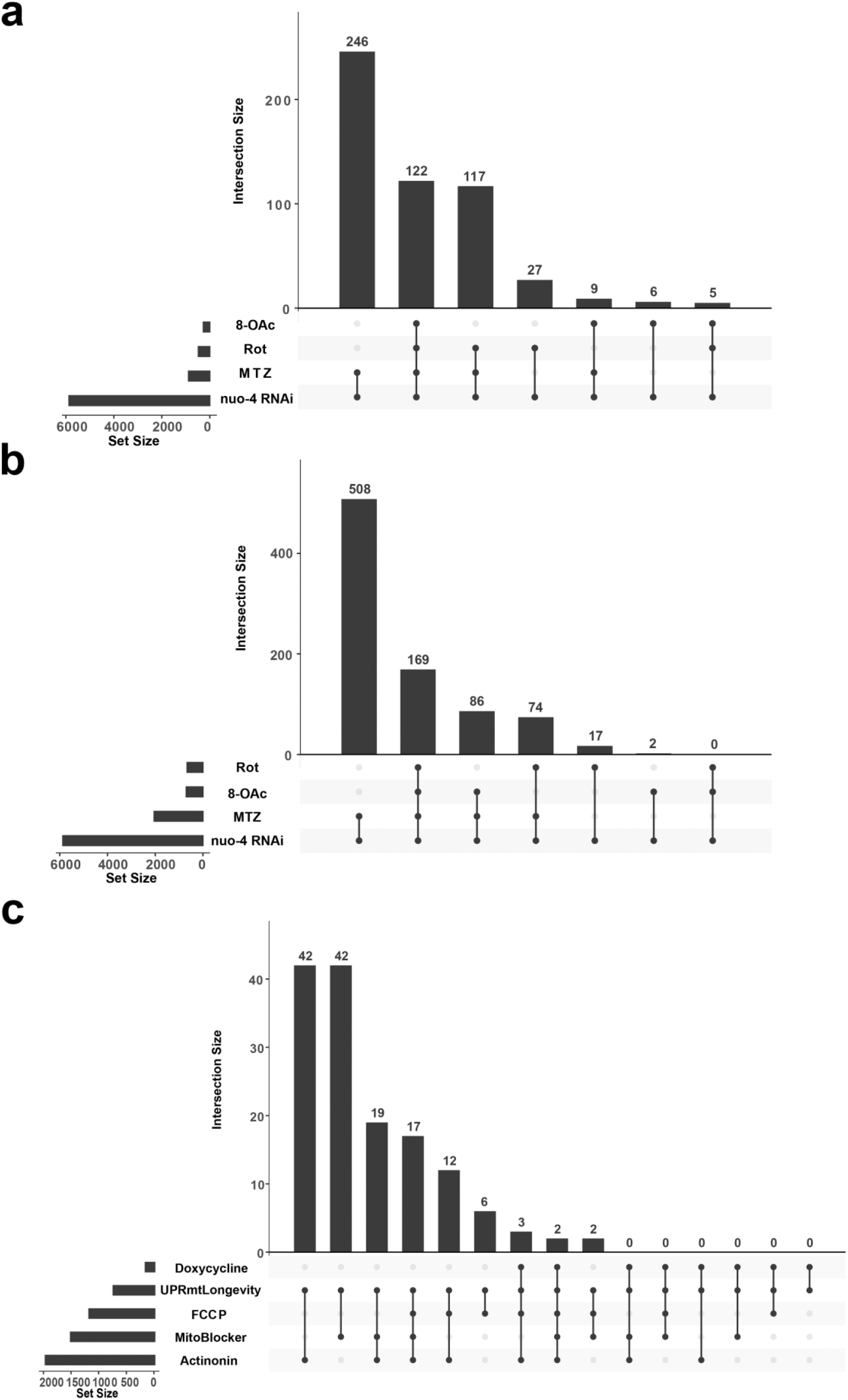
Comparison of UPR^mt^ induction transcriptome across species. **(a)** Upset plot comparing the UPR^mt^ DEG sets in *C. elegans* induced by *nuo-4* RNAi to chemical treatments (mycothiazole (MTZ), 8-O-acetylmycothiazole (8-OAc), and rotenone (Rot)). **(b)**Upset plot comparing the UPR^mt^ DEG sets in *C. elegans* induced by *nuo-4* RNAi to the UPR^mt^ orthologous DEG sets in Huh7 liver carcinoma cells treated with MTZ, 8-OAc, and Rot. **(c)** Upset plot comparing our novel UPR^mt^ driven longevity DEG set (UPRmtLongevity) to the DEG set of the mammalian integrated stress response (ISR) from 293T cells. All the DEGs were determined by adj p-val < 0.05.

Next, we show examples of how our genesets can be compared to non-*C. elegans* data by comparing our dataset to human mitochondrial stress conditions. First, we compare our *nuo-4* RNAi condition to the transcriptome of human cancer cell (Huh7 cell) treated with the same chemicals: MTZ, 8-OAc, and rotenone ^44^. 169 shared DEGs were identified between our UPR^mt^ DEG set and the chemical-induced UPR^mt^ transcriptome from Huh-7 cell line (**Fig. 6b**). Finally, we compared our longevity associated UPR^mt^ driven longevity DEGs (UPRmtLongevity) that were upregulated in all the long-lived UPR^mt^ activations (knocked down with *atp-2*, *cco-1*, *cyc-1*, *nuo-4 and mrps-5*) and downregulated in the short-lived UPR^mt^ activations (knocked down with *mev-1*, *fzo-1* and *tomm-20*) with the transcriptome of ISR activated non-cancer human cell lines. The UPRmtLongevitygeneset consisted of total 759 DEGs (525 upregulated DEGs and 234 downregulated DEGs) and was compared with the publicly available data (GSE84631) of the RNA sequencing data of ISR from 293T cells ^45^ following the ortholog conversion using OrthoList2. We identified 17 overlapping DEGs in all the conditions excluding Doxycycline, presumably due to the small set size of the DEGs from Doxycycline treated cells. There were 42 shared DEGs between our UPRmtLongevity geneset and Actinonin treated cells, and 42 shared DEGs between UPRmtLongevity geneset and MitoBlocker treated cells (**Fig. 6C**). Although there were some overlapping DEGs, the lack of more robust overlap does potentially suggest that *C. elegans* UPRmt and mammalian mitochondrial stress response are in fact distinct, though further directed studies will need to be performed to make conclusions. Overall, our UPR^mt^ DEG set can robustly be used to identify UPR^mt^ in *C. elegans* upon different activation and can further be used to compare differences in UPR^mt^ across organisms.

### Usage Notes

Overall, the data presented here offer a comprehensive analysis of diverse UPR^mt^ activation conditions in *C. elegans*. These carefully controlled datasets allow researchers to identify distinct and shared gene expression changes across distinct mitochondrial stress conditions. Moreover, these data can be easily compared to a researcher’s own dataset to identify shared or distinct gene expression changes for other mitochondria-, stress-, and age-associated interventions.

The readers can download the dataset to find the DEGs using different normalization methods or significance cutoff (See **Data Records**). The usage of our new UPR^mt^ genesets proposed in **Table S1** (525 upregulated, 234 downregulated DEGs) is recommended for assessment of UPR^mt^ activation in longevity studies of *C. elegans* as shown in the example of **Fig. 6c**. The genes indicated in **Fig. 3e** are those that are specifically upregulated upon UPR^mt^ activation associated with longevity, and thus are better candidates for single-gene reporters to measure UPR^mt^ activation associated with longevity. Furthermore, the DEGs commonly up- or downregulated after each ETC knockdown will provide useful information on the genes regulated after activation of UPR^mt^ by ETC perturbation.

## Code Availability

All R codes used in this study are uploaded in the Annotare accession E-MTAB-15031 along with the raw data as a downloadable txt file.

## Acknowledgements

J.K. is supported by the USC Provost Fellowship; G.G. is supported by T32AG052374 and R01AG079806-02S1 from the National Institute on Aging; and R.H.S. is supported by R01AG079806 from the National Institute on Aging and 2022-A-010-SUP from the Larry L. Hillblom Foundation. Some strains were provided by the CGC, which is funded by the NIH Office of Research Infrastructure Programs grant P40 OD010440. Some gene analysis was performed using Wormbase, which is funded on a U41 grant HG002223. We thank Dr. Bérénice Benayoun and the members of the Benayoun lab for availability of code and assistance with RNA-seq analysis.

## Author contributions

J.K. and R.H.S conceptualized the study. J.K. conducted RNA-seq analysis, visualized the data, and wrote the manuscript. J.K., N.D., and R.H.S. performed the *C. elegans* experiments. G.G. assisted in the RNA-seq analysis, supporting data processing. All authors reviewed, edited, and approved the final manuscript.

## Figure Legends

**Supplementary Figure 1.**
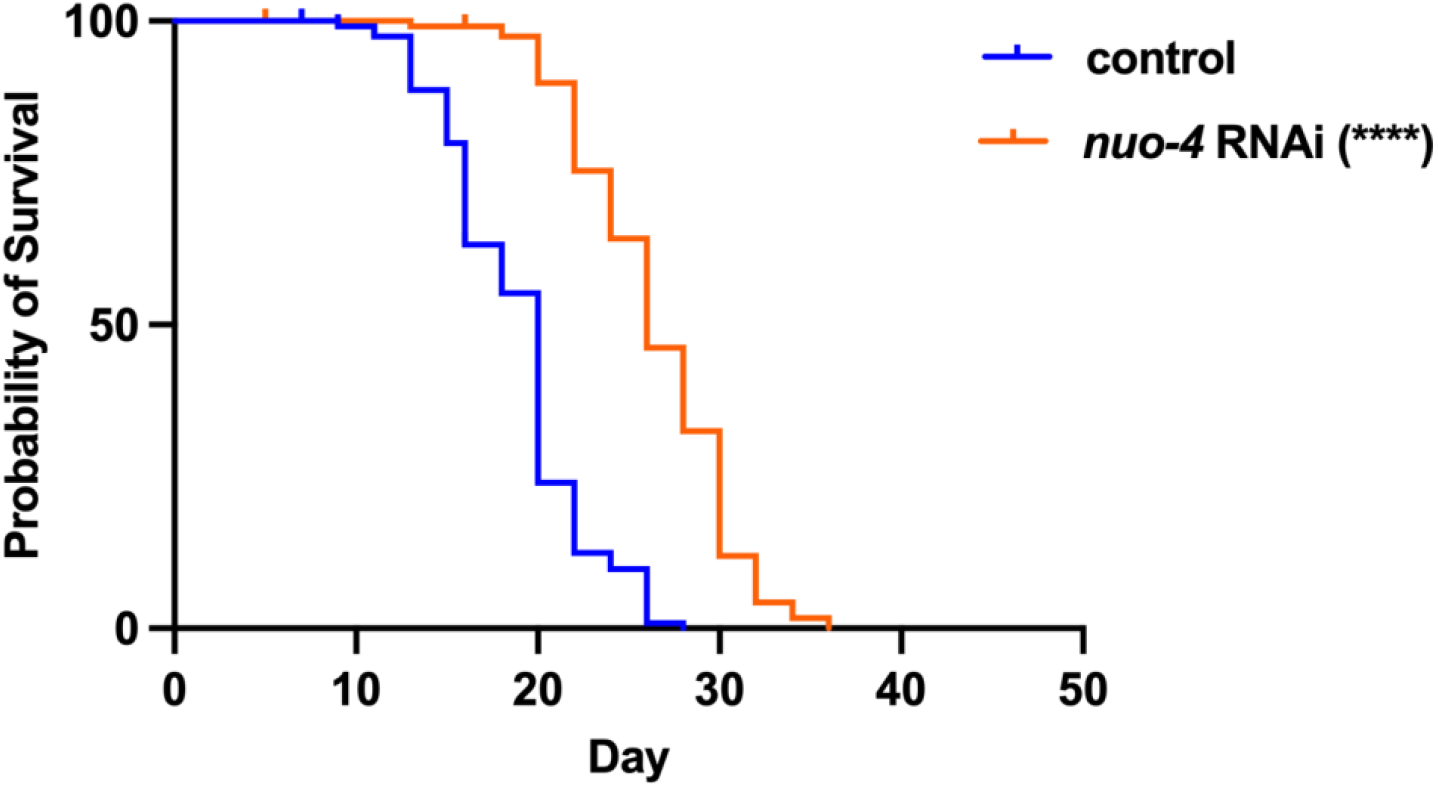
Survival curve of *nuo-4* RNAi treated worms. nuo-4 knockdown worms are longer-lived (N=117; median lifespan of 26 days) compared to the control group (N=113; median lifespan of 20 days). p-value < 0.0001. Data is representative of 3 independent trials.

## Tables

**Table 1. Respective RNAi knockdown targets and lifespans Supplementary Table 1. QC analysis of RNA.**

**Supplementary Table 1. QC analysis of RNA.**

**Supplementary Table 2. Comprehensive upregulated DEGs that are commonly expressed in long-lived animals but not in short-lived animals.**

**Supplementary Table 3. Comprehensive downregulated DEGs that are commonly expressed in long-lived animals but not in short-lived animals.**

